# Impact of acute visual experience on development of LGN receptive fields in the ferret

**DOI:** 10.1101/2021.07.16.452697

**Authors:** Andrea K. Stacy, Nathan A. Schneider, Noah K. Gilman, Stephen D. Van Hooser

## Abstract

Selectivity for direction of motion is a key feature of primary visual cortical neurons. Visual experience is required for direction selectivity in carnivore and primate visual cortex, but the circuit mechanisms of its formation remain incompletely understood. Here we examined how developing lateral geniculate nucleus (LGN) neurons may contribute to cortical direction selectivity. Using *in vivo* electrophysiology techniques, we examined LGN receptive field properties of visually naïve female ferrets before and after exposure to 6 hours of motion stimuli in order to assess the effect of acute visual experience on LGN cell development. We found that acute experience with motion stimuli did not significantly affect the weak orientation or direction selectivity of LGN neurons. In addition, we found that neither latency nor sustainedness or transience of LGN neurons significantly changed with acute experience. These results suggest that the direction selectivity that emerges in cortex after acute experience is computed in cortex and cannot be explained by changes in LGN cells.

**Significance Statement:** The development of typical neural circuitry requires experience-independent and experience-dependent factors. In the visual cortex of carnivores and primates, selectivity for motion arises as a result of experience, but we do not understand whether the major brain area that sits between the retina and the visual cortex – the lateral geniculate nucleus of the thalamus – also participates. Here we found that lateral geniculate neurons do not exhibit changes as a result of several hours of visual experience with moving stimuli, at a time when visual cortical neurons undergo a rapid change. We conclude that lateral geniculate neurons do not participate in this plasticity, and that changes in cortex are likely responsible for the development of direction selectivity in carnivores and primates.

## Introduction

Detection of direction of motion is a key component of mammalian visual processing that emerges early in the visual pathway. In primates and carnivores, direction selectivity is thought to first emerge in the primary visual cortex (V1) (Hubel and Wiesel, 1962; Livingstone, 1998), and requires visual experience for its development (Chapman and Stryker, 1993; Horton and Hocking, 1996; White et al., 2001; Li et al., 2006). Early deprivation of visual experience in kittens (Zhou et al., 1995) and ferrets (Li et al., 2006) impairs development of direction selectivity, and this deficit cannot be recovered despite subsequent introduction to normal visual experience (Li et al., 2006). However, direction selectivity can be rapidly and artificially induced in visually naïve ferrets with only 3-6 hours of visual exposure to moving stimuli (Li et al., 2008; Van Hooser et al., 2012; Ritter et al., 2017) allowing us to probe its development in a laboratory setting to study its mechanisms.

What circuitry underlies this rapid developmental emergence? The major feed-forward input to the visual cortex arrives via the lateral geniculate nucleus (LGN), and here we sought to understand what, if any, contribution to emerging direction selectivity was made by changes in LGN neurons. While the developing visual cortex exhibits rapid changes in direction selectivity with a few hours of visual experience, it is unknown if developing LGN cells also exhibit substantial receptive field plasticity. One could imagine two major hypotheses for how potential developmental plasticity in LGN neurons might influence direction selectivity in V1.

First, LGN neurons could acquire some substantial direction selectivity themselves as a result of experience, so that the input that each cell provides to cortex exhibits increased direction selectivity as a result of experience. While this hypothesis would be at odds with the “spot detector” model for a majority of LGN receptive fields (Kuffler, 1953) recent studies have shown that some species have substantial direction-selective or orientation-selective channels that run through the LGN (Marshel et al., 2012; Cruz-Martin et al., 2014; Sun et al., 2016; Hillier et al., 2017) so this is an important possibility to assess.

Second, LGN cells themselves may be unselective for orientation or direction, but there could be experience-dependent changes in their receptive fields that are necessary for the expression of direction selectivity in visual cortex. For example, direction selectivity requires some spatiotemporal offset in inputs (Reichardt, 1961; Barlow and Levick, 1965; Adelson and Bergen, 1985; Watson and Ahumada, 1985; Suarez et al., 1995; Maex and Orban, 1996) such that inputs to one side of the cell’s receptive field have different latencies than the inputs to the other side (Movshon et al., 1978a; McLean and Palmer, 1989; Albrecht and Geisler, 1991; Reid et al., 1991; Saul and Humphrey, 1992; DeAngelis et al., 1993; Jagadeesh et al., 1997; Livingstone, 1998; Priebe and Ferster, 2005). Perhaps visual experience is required to establish some property, such as a spread in LGN cell latencies (Saul and Humphrey, 1990, 1992; Ferster et al., 1996; Alonso et al., 2001; Stanley et al., 2012) or a change in the sustained/transient nature of responses (Marr and Ullman, 1981; Lien and Scanziani, 2018) that is necessary for direction selectivity to be computed in cortex, even if the computation is carried out in cortex.

Using *in vivo* electrophysiology, we examined the receptive field properties of individual thalamic neurons before and after acute visual stimulation with drifting sinusoidal gratings. We found that orientation selectivity, direction selectivity, response latency, and sustainedness/transience of individual LGN neurons did not significantly change with acute visual experience. That is, we found no evidence of rapid changes in LGN receptive fields when cortex is undergoing a substantial increase in direction selectivity. These results are consistent with the idea that the changes driving an increase in cortical direction selectivity are first occurring in V1, either at the convergence of LGN inputs or in the V1 circuitry itself.

## Materials and Methods

### EXPERIMENTAL DESIGN

All experimental procedures were approved by the Brandeis University Animal Care and Use Committee and performed in compliance with National Institutes of Health guidelines. Ferrets (*Mustela putorius furo*; n = 29), age postnatal day 30-35 (P30–35) were used for electrophysiological experiments. Females were used because young animals required cohousing with sexually mature females, and cohousing with males causes undue stress in mature females.

### SURGICAL PROCEDURES

Ferrets were initially anesthetized with ketamine (20mg/kg i.m.). Atropine (0.16-0.8 mg/kg i.m.) and dexamethasone (0.5 mg/kg i.m.) were used to reduce salivary and bronchial secretion and to reduce inflammation and swelling, respectively. The animal was anesthetized with a mixture of isoflurane, oxygen, and nitrous oxide through a mask while a tracheostomy was performed. After completion of the tracheostomy, animals were ventilated with 1-2% isoflurane in a 2:1 mixture of nitrous oxide and oxygen. Next, a cannula was inserted into the intraperitoneal cavity for delivery of 5% dextrose in lactated Ringer’s solution (3 ml/kg/h). Body temperature was maintained at 37°C using a thermostatically controlled heating pad. End-tidal CO_2_ levels and respiration rate were monitored and kept within the appropriate physiological range (3.5–4%). The animal was held in place by a custom stereotaxic frame that did not obstruct vision. Silicone oil was placed on the eyes to prevent corneal damage. An O-ring (10mm diameter, 2mm thickness) was cemented to the skull using Zap-It gel and zip kicker over LGN in the right hemisphere, and a 6 × 6 mm craniotomy was made in the interior of the O-ring. The dura was removed with a 31-gauge needle. All wound margins were infused with bupivacaine. Before beginning electrophysiological recordings, the ferret was paralyzed with the neuromuscular blocker gallamine triethiodide (10–30 mg/kg^/^h) through the intraperitoneal cavity in order to suppress spontaneous eye movements, and the nitrous oxide-oxygen mixture was adjusted to 1:1. The animal’s ECG was continuously monitored to ensure adequate anesthesia, and the percentage of isoflurane was increased if the ECG indicated any distress.

### ELECTROPHYSIOLOGY

Following removal of the dura, warm agarose (2-4% in phosphate-buffered saline) was applied to the craniotomy to prevent brain pulsation. Then, a custom (https://github.com/VH-Lab/vhlab-parts) 3D-printed chamber-and-grid (Realize Inc.) was cemented to the O-ring using Zap gel (Zap Adhesives) and Zip Kicker (ZAP) so that an electrode could be driven through the grid and inserted perpendicularly to the surface of the brain. A low-impedance tungsten electrode (World Precision Instruments, 0.1MΩ, part TM33B01) was inserted through different grid holes using an MP-285 manipulator (Sutter Instruments) until the LGN was initially located by monitoring modulation to handheld stimuli on a loudspeaker. The LGN was then mapped by driving the electrode through adjacent holes and a recording location was identified using a signal-to-noise ratio sufficient for isolation as criteria. Then, either a linear microelectrode array, microwire brush array (Microprobes), or a tetrode array (Plexon S-probe) was inserted to the same location for recording. The signal was amplified using an RHD2000 amplifying/digitizing chip and USB interface board (Intan Technologies) and stimulus timing information acquired using a Micro1401 acquisition board and Spike2 software (Cambridge Electronic Design). Units were initially automatically sorted offline by a mixture-of-Gaussians model using the KlustaKwik algorithm (http://klustakwik.sourceforge.net; Harris et al., 2000), and manually adjusted using custom software in MATLAB. Recordings were made at the beginning of the experiment and after 6 hours of training with visual stimuli.

### HISTOLOGY

Upon completion of experiments, an electrode coated in the fluorescent dye DiI (DiCarlo et al., 1996) was inserted at the same grid hole location and depth and left in place for 20 minutes. Animals were then transcardially perfused and the brain was placed in 4% paraformaldehyde in 0.1M PBS at 4°C for 24 hours and then moved to 10% sucrose in PBS for 24-48 h. This was followed by placement in 30% sucrose in PBS at 4°C until sectioning. The recording hemisphere of the brain was sectioned sagittally into 50-60 μm sections using a sliding microtome (Leica SM2010R). All staining procedures were performed on a shaker. We washed sections in 0.1M PBS 3×5 min and permeabilized in 0.3% Triton-X 100 diluted in PBS for 2 hours at room temperature. Then slices were incubated in fluorophore-conjugated anti-NeuN antibody (Alexa Fluor 488 Rabbit anti NeuN, Millipore ABN78A4) at a 1:300 dilution overnight (>12 hours) at 4°C. Sections were then washed 3×5 min in PBS and mounted on slides until air-dried. Slides were then cover-slipped with Fluoromount-G media (Electron Microscopy Sciences, Ft. Washington, PA) and edges were sealed using nail polish (Electron Microscopy Sciences, Ft. Washington, PA). Histological sections were viewed using a fluorescent microscope (Keyence BX-Z 710) and electrode tracks were reconstructed using DiI dye traces.

### VISUAL STIMULATION

Visual stimuli were created in MATLAB (MathWorks) using the Psychophysics Toolbox (Brainard, 1997; Pelli, 1997) and displayed on a 21-inch flat face CRT monitor (GDM-520, Sony) with a resolution of 800 × 600 and a refresh rate of 100 Hz. The monitor was placed at a distance such that it subtended 51 × 42 degrees of visual space. We manually mapped receptive fields by displaying circular patches of drifting sinusoidal gratings at different positions and moving the monitor to accommodate different eccentricities while listening to the responses on a loudspeaker.

Drifting grating stimuli were full-field, high-contrast sinusoidal gratings (2 s duration; 3.5 s interstimulus interval) presented pseudorandomly, with direction of motion (in steps of 45°) in either of the two directions orthogonal to the axis of orientation. For spatial and temporal frequency tuning, direction was co-varied with stimuli consisting of drifting sinusoidal gratings at 6 different spatial frequencies (0.01-0.32 cpd) and 6 different temporal frequencies (0.5-32 Hz), respectively at 100% contrast. Spatial phase stimuli consisted of counterphase static sinusoidal gratings (spatial frequency 0.08 cycles per degree and temporal frequency 1Hz) at six different spatial phases.

### DATA ANALYSIS

We recorded from a total of 543 LGN neurons as identified by spike sorting. For drifting gratings that examined orientation/direction, we examined mean responses (F0) and modulation at the stimulus frequency (F1). If a cell’s F1 response was greater than the mean response (F0), F1 was used to calculate index values. If F0 was greater, it was used for calculations (Movshon et al., 1978a, b; Heimel et al., 2005). Neurons were included in analysis if they exhibited significant variation across all conditions by an ANOVA test, p<0.05. We calculated responses for 207 neurons. The fitless vector measures, circular variance (CV) and directional circular variance (DCV), were calculated as previously described (Ringach et al., 2002; Mazurek et al., 2014) and 1-CV and 1-DCV were used to quantify selectivity for orientation and direction, respectively.

For spatial phase stimuli, the spatial phase that elicited the greatest response (F1) was chosen for further analysis. Raster responses from 200 trials were averaged for peristimulus time histograms (PSTH; 10-ms time bins) of cell responses and PSTHs were used to calculate peak rate, maintained rate, response latency and transience/sustainedness. Values were calculated as previously described (Van Hooser et al., 2003). In brief, responses were binned into 10-ms increments. The bin with the greatest number of spikes was used for peak latency. Peak firing rate (PR) was defined as the mean rate during the 10-ms bin centered on the peak latency. Initial latency was taken as the first bin with spike count greater than or equal to one half the maximum. There was one cell that responded with such a delayed latency that its peak response occurred in the opposite phase. We assigned the latency of this cell to be 500ms + peak rate. Maintained firing rate (MR) was defined as the mean firing rate during a 100-500 ms window after the peak latency time. Transience was defined as the transient time constant, T_trans_, where MR = PR * exp(−300 ms/T_trans_).

## Results

At the time of eye-opening in ferrets, corresponding to approximately postnatal age 30-33 (P30-33), neurons in the primary visual cortex (V1) are selective for orientation of stimuli but are not selective for direction of motion of stimuli (Li et al., 2006; Li et al., 2008). Direction selectivity develops with a few days of visual experience and increases to mature levels within the first 2-3 weeks of visual experience (Li et al., 2006). Furthermore, experience is required for the increase in cortical direction selectivity. Animals that have been dark-reared at the time of eye-opening until P45 never develop cortical selectivity for stimulus direction even if they are typically reared beginning at P45 (Li et al., 2006). However, direction selectivity can be rapidly induced in a laboratory environment by providing young ferrets with 3-6 hours of exposure to drifting gratings (Li et al., 2008; Van Hooser et al., 2012).

In order to understand how the receptive field properties of LGN cells may be changing during this time of great plasticity of receptive fields in visual cortex, we made *in vivo* electrophysiological recordings in the LGN of visually naïve ferrets (P30-35; n=29). To measure the responses of cells in the naïve state, we needed to use methods that would allow us to record the responses of many LGN neurons simultaneously. The traditional single cell recording approach of carefully introducing stimuli to an individual neuron for an hour or more would have meant that the third or fourth neuron recorded would have undergone extensive visual experience. The ferret LGN is small and is located approximately 1 cm below the brain surface, so we developed a grid-and-chamber system (**Figure 1a**) that allowed us to map the LGN with low impedance electrodes before precisely introducing a multi-channel recording electrode to the brain at the same insertion position and direction as our preferred mapping penetration (**Figure 1b**).

**Figure 1.**
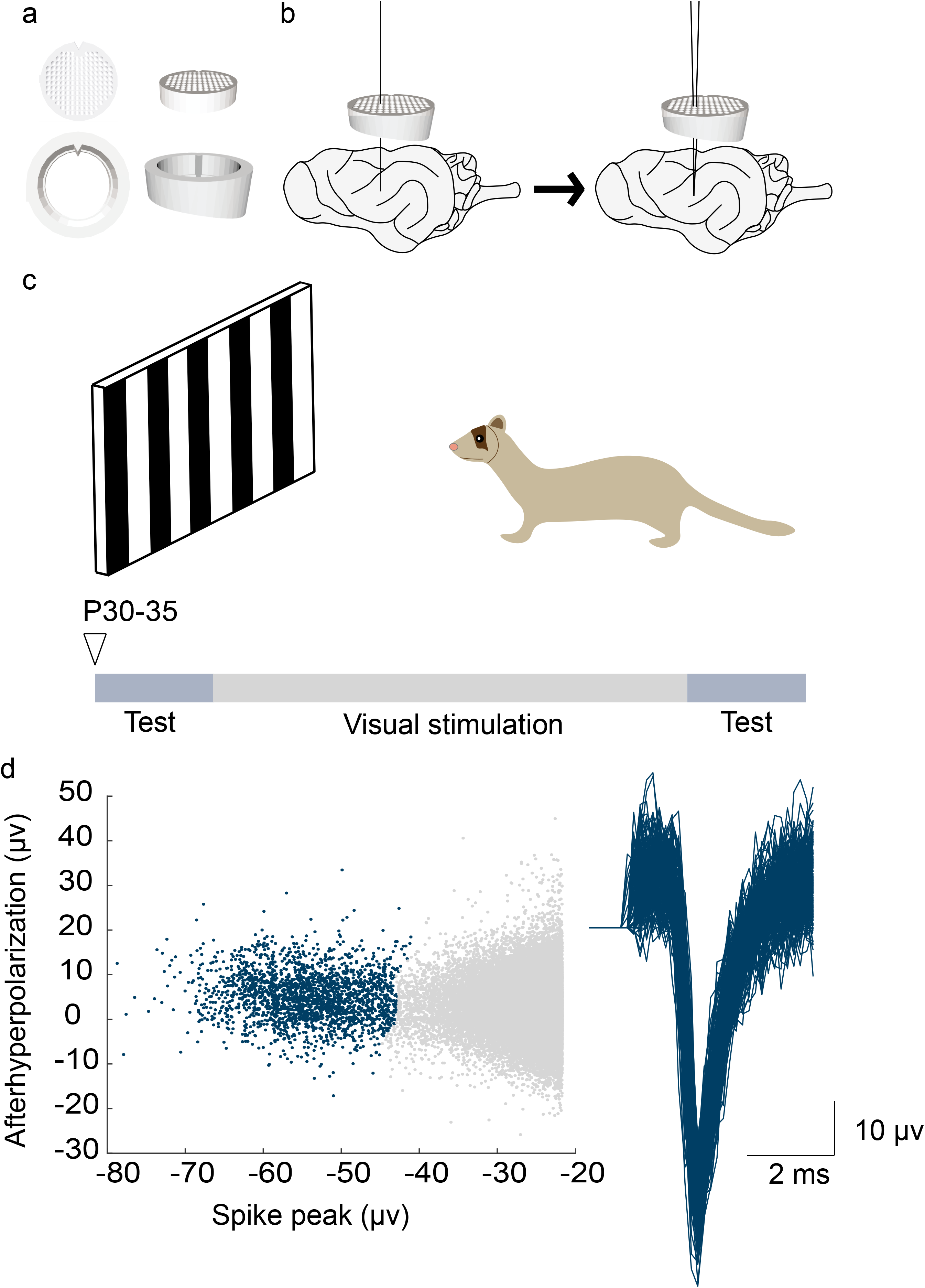
Methods for single-channel mapping and multi-channel recording from ferret lateral geniculate nucleus (LGN). **a**. Top-down and side-view of grid-and-chamber design used for locating and mapping LGN. **b**. Schematic for initial mapping and placement of multi-channel recording electrode for experiment duration. Grid-and-chamber was secured to the skull of the animal and a single-channel mapping electrode was lowered into different grid holes and responses to a light stimulus were monitored. When visual responses characteristic of LGN were located, consecutive grid holes were mapped with the single-channel electrode until a location with a central receptive field with high signal-to-noise ratio was identified. Then, the multi-channel electrode was lowered into the same location for the experiment. **c**. Experimental setup with timeline of experimental paradigm. Experiment began with a test phase to record initial responses to moving sinusoidal gratings of varying orientations, directions, spatial and temporal frequencies, and spatial phases. Then, the animal was exposed to six hours of visual stimulation with bi-directional sinusoidal gratings moving at 0 or 90 degrees. Following this, the same set of test stimuli were used to measure responses again. **d**. Stably isolated spike cluster and waveform recorded from a neuron across test phases.

After making initial receptive field measurements, we provided the animal with 6 hours of visual stimulus exposure to a “training” motion stimulus that consisted of drifting sinusoidal gratings (0.08 cpd; temporal frequency 1-4 Hz) moving back and forth along the axis of motion orthogonal to the grating orientation (Li et al., 2008; Van Hooser et al., 2012; Ritter et al., 2017; Roy et al., 2020) (**Figure 1c**). In this way, we obtained measures of LGN receptive fields “before” and “after” visual experience. In this work, while we were able to isolate single units (**Figure 1d**), we did not find that we could commonly hold the same cells before and after visual experience, so we analyzed these groups of neurons as separate populations and intended no claims about following receptive fields of individual neurons, as the lab has previously done in 2-photon imaging work (Li et al., 2008; Van Hooser et al., 2012; Roy et al., 2020).

### Orientation and direction selectivity

We wanted to test the hypothesis that acute visual experience might cause an increase in orientation or direction selectivity in individual LGN cells. If this were true, then the rapid increase in direction selectivity that is observed in visual cortical neurons might reflect a direct contribution of individual LGN inputs.

The strengths of orientation and direction tuning were quantified by calculation of an orientation selectivity index (1-CV; see Materials and Methods) and a direction selectivity index (1-DirCV; see Materials and Methods).

We first compared the orientation selectivity values of naïve LGN cells to a population of naïve and experienced ferret visual cortical cells that were recorded with the same stimuli, experimental setup, and analysis code and reported in Clemens et al. (2012). Initial orientation indexes for LGN cells were weak, exhibiting a median value of 0.12. Representative tuning curves for naïve LGN cells are shown in **Figure 2a** (top row) and tuning curves for naïve and experienced cortical cells from layer II-VI are shown in **Figure 2a,b** (bottom rows). We found that orientation selectivity index values in naïve LGN neurons (median: 0.12) were much lower than in naïve (median: 0.435; Kruskal-Wallis test H(1)=87.2433, p=0) and experienced (median: 0.51; Kruskal-Wallis test H(1)=156.2993, p=0) cortical neurons (**Figure 2c, left**).

**Figure 2.**
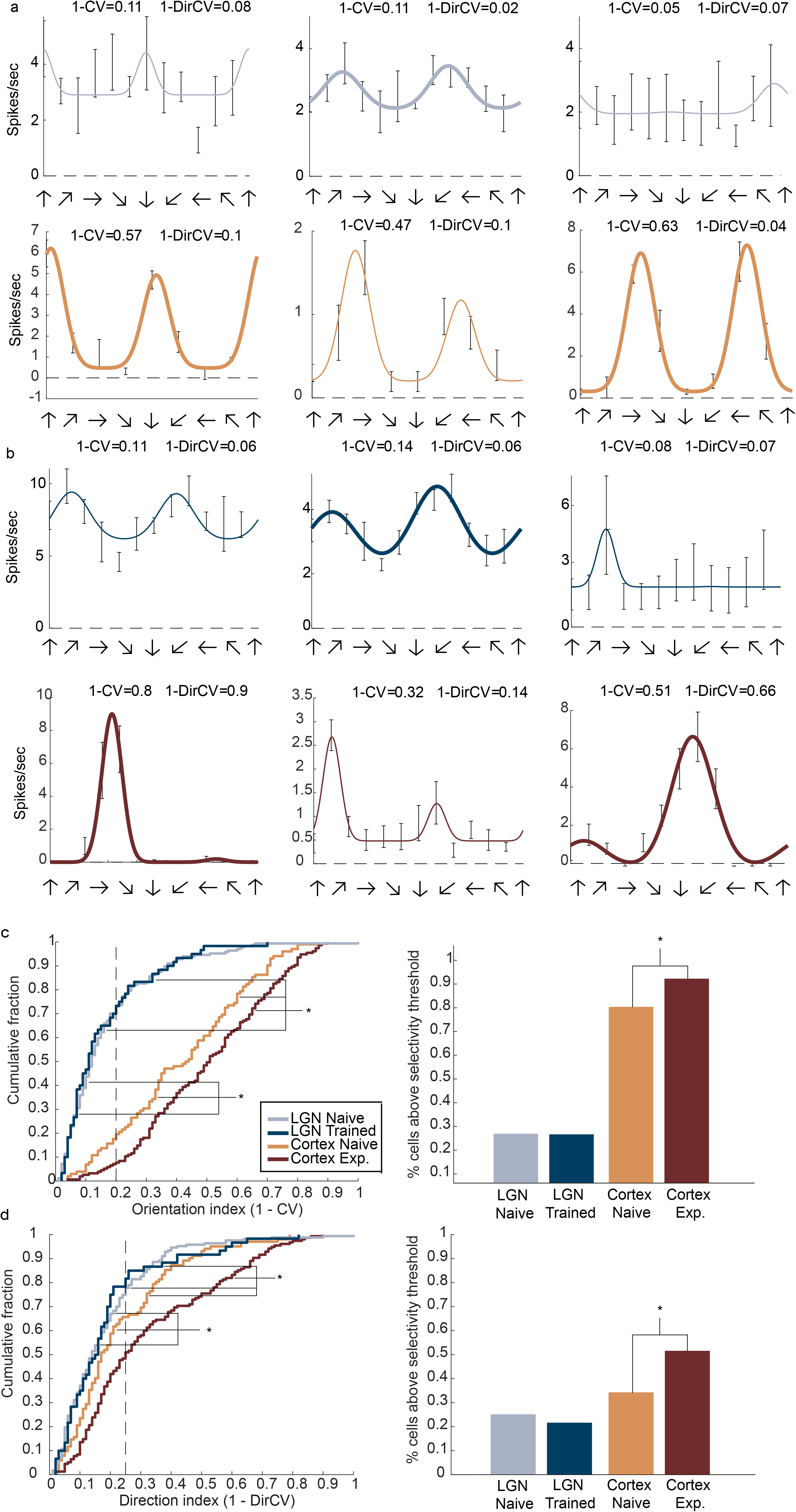
Lack of increase in orientation and direction selectivity with 6 hours of experience with moving stimuli. **a**. Tuning curves of 3 example LGN neurons (top row) before visual stimulation and 3 naïve primary visual cortex neurons (V1; bottom row; Clemens et al., 2012). Bolded fit shows significant orientation tuning (Hotelling-T2, p<0.05), non-bolded fits not significant. Dashed line indicates response to control gray screen. **b**. Tuning curves of 3 example LGN neurons (top row) in ferrets after visual stimulation and 3 experienced V1 neurons (bottom row). Conventions as in Figure 2a. **c (left).** Cumulative histogram of orientation selectivity in LGN before (LGN Naïve, n=167) and after (LGN Trained, n=60) 6 hours of visual stimulation compared to data previously collected from immature (PND<35, n=102) and experienced (PND>35, n=155) ferret primary visual cortex (V1) (Clemens et al., 2012). Degree of orientation selectivity is expressed as 1-CV in orientation space, n indicates cell number, * indicates reference group, and lines point to groups that are significantly different from the reference group. Orientation selectivity index values in naïve LGN neurons (median: 0.12) were much lower than in naïve (median: 0.435; Kruskal-Wallis test H(1)=87.2433, p=0) and experienced (median: 0.51; Kruskal-Wallis test H(1)=156.2993, p=0) cortical neurons. Visually naïve LGN neurons did not significantly increase in their selectivity for orientation following 6 hours of training (median: 0.11; Kruskal-Wallis test, H(1)=0.1037, p=0.7474). Dotted line indicates arbitrary threshold for marginal selectivity. **c (right).** Percentage of cells that surpass arbitrary threshold for marginal orientation selectivity for each group. There was no significant difference in proportion of the orientation-selective cell populations in LGN before and after training (x^2^(1)=0.0018, p=0.9666). Proportion of orientation-selective cells in cortex increased significantly (x^2^(1)=6.9206, p=0.0085). **d (left).** Cumulative histogram of direction selectivity in LGN before (LGN Naïve) and after (LGN Trained) 6 hours of visual stimulation compared to data previously collected from immature (PND<35) and experienced (PND>35) ferret V1 (Clemens et al., 2012). Degree of direction selectivity is expressed as 1-DirCV in direction space. Conventions as in Figure 2c. Direction selectivity index values in naïve LGN neurons (median: 0.14) were lower than in either naïve (median: 0.17: Kruskal-Wallis test H(1)=6.8602, p=0.0088) or experienced (median: 0.25.; Kruskal-Wallis test H(1)=42.2017, p=8.2329e-11) cortical neurons. Visually naïve LGN neurons did not significantly increase in their selectivity for direction following 6 hours of training (median: 0.155; Kruskal-Wallis test, H(1)=0.0012, p=0.9726). **d (right).** Percentage of cells that surpass arbitrary threshold for marginal direction selectivity for each group. There was no significant difference in proportion of direction-selective cells before and after training ( x^2^(1)=0.2917, p=0.5891). Proprotion of direction-selective cells in cortex significantly increased with experience (x^2^(1)=7.4461, p=0.0064).

We next asked whether 6 hours of visual experience, which causes rapid and reliable induction of direction selectivity in visual cortex (Li et al., 2008; Van Hooser et al., 2012; Roy et al., 2016; Ritter et al., 2017; Roy et al., 2020), caused changes in the amount of orientation selectivity measured in LGN cells. We found that visually naïve LGN neurons did not significantly increase their selectivity for orientation following 6 hours of training (median: 0.11; Kruskal-Wallis test, H(1)=0.1037, p=0.7474; (**Figure 2c, left**).

We then compared the LGN orientation tuning values for the after condition to the naïve and experienced cortical cells. Representative tuning curves for trained LGN cells and experienced V1 cells are shown in **Figure 2a and b**, bottom rows, respectively. We found that LGN orientation tuning values for the after condition were significantly lower than the values of the visually naïve (Kruskal-Wallis test, H(1)=55.270 p=1.0503e-13) and experienced V1 neurons (Kruskal-Wallis test, H(1)=87.1024 p=0). When we compared orientation-selectivity values for visually naïve LGN neurons to LGN neurons following 6 hours of training, we found no significant difference (Kruskal-Wallis test, H(1)=0.1037, p=0.7474). Further, when we compared the proportion of the orientation-selective cell populations in LGN before and after training, we found no significant differences (x^2^(1)=0.0018, p=0.9666). However, when we compared the orientation-selective proportions of V1 cells we found a significant increase (x^2^(1)=6.9206, p=0.0085) (**Figure 2c, right**).

After examining orientation index values, we compared direction selectivity index values of naïve LGN neurons (median: 0.14) to V1 values and found that they were lower than naïve (median: 0.17: Kruskal-Wallis test H(1)=6.8602, p=0.0088) and experienced (median: 0.25.; Kruskal-Wallis test H(1)=42.2017, p=8.2329e-11) cortical neurons (**Figure 2d, left**). We then compared direction-selectivity values for trained LGN neurons to naïve and experienced V1 cells and found no significant differences (Kruskal-Wallis test, H(1)=3.9565 p=0.0467; Kruskal-Wallis test, H(1)=22.4016, p=2.2119e-6). When we compared direction-selectivity values for visually naïve LGN neurons to LGN neurons following training, we found no significant difference (median: 0.155; Kruskal-Wallis test, H(1)=0.0012, p=0.9726) (**Figure 2d, left**). Additionally, when we compared the proportion of direction-selective cell populations in LGN before and after training we found no significant difference (x^2^(1)=0.2917, p=0.5891). However, when we compared the direction-selective population proportions before and after experience in cortical cells we found a significant increase (x^2^(1)=7.4461, p=0.0064) (**Figure 2d, right**).

### Sustainedness/Transience

Our first analyses showed that LGN cells did not exhibit increases in orientation or direction selectivity themselves, but it remained possible that other short-term changes in receptive field properties could aid a cortical calculation of direction selectivity. The circuitry that underlies direction selectivity in carnivore visual cortex is still a matter of research, but we can gain some clues about what receptive field properties might be important for the input to such a circuit from the literature.

Marr and Ullman hypothesized that direction selectivity might arise by combining a low-latency, transient input at one location (such as a Y cell) with a longer-latency, sustained input (such as an X cell) at a neighboring location to create a direction-selective cell (Marr and Ullman, 1981). This neuron would be selective for motion that arose on the side of the sustained input’s location and moved in the direction of the transient input (**Figure 3a**). Evidence for this arrangement was found in the projections of the mouse LGN to direction-selective cells in visual cortex (Lien and Scanziani, 2018). If there were a major experience-dependent change in the sustainedness or transience of LGN cells, such as the appearance of sustained responses or transient responses, then the availability of these signals could impact the calculation of cortical direction selectivity.

**Figure 3.**
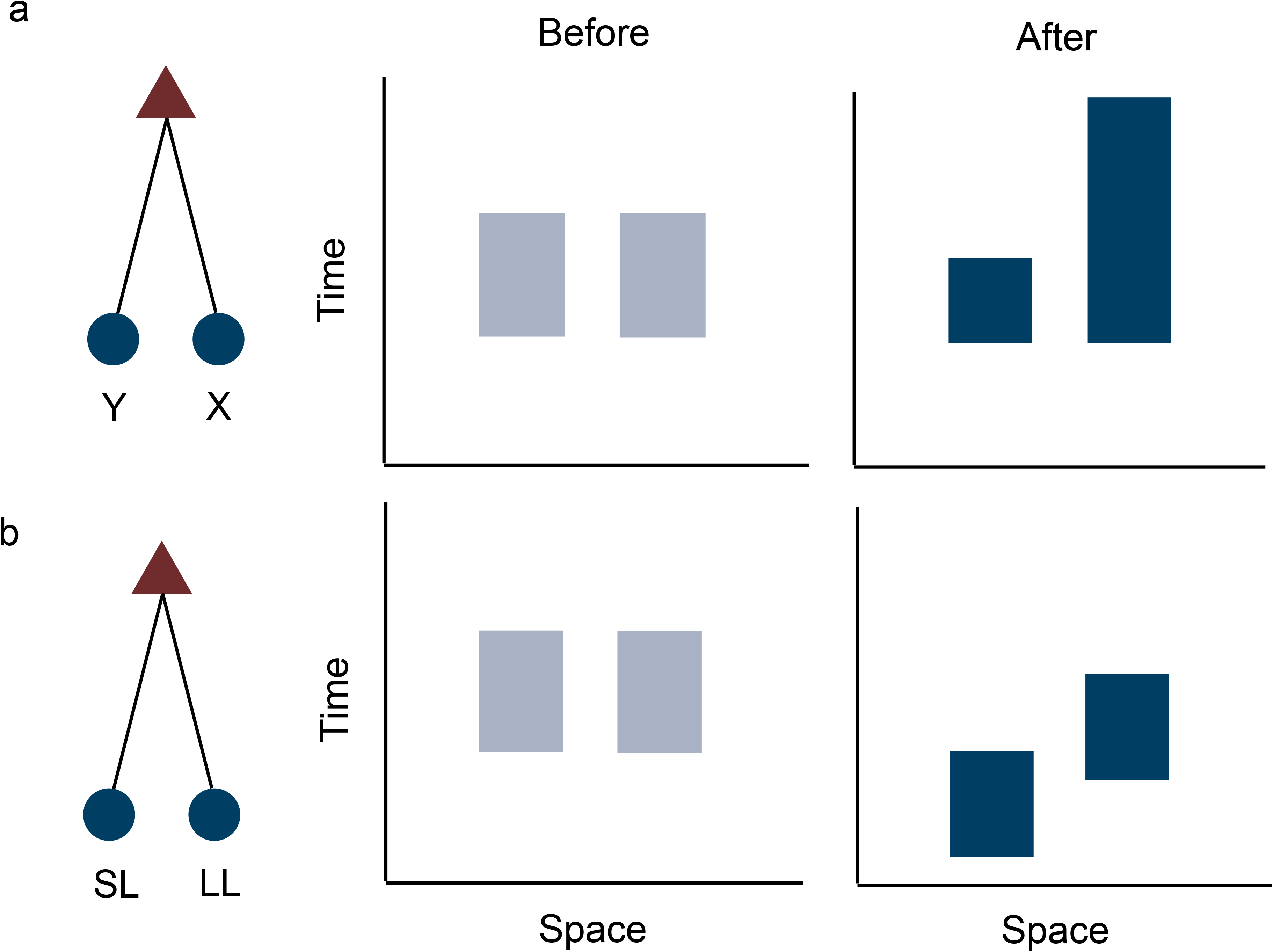
Schematic illustrating how direction selectivity could arise in V1 from combined LGN inputs. **a.** A direction-selective V1 cell is created by combining an input at one location (such as a Y cell) with a sustained input (such as an X cell) at a neighboring location to create a direction-selective cell. Hypothesized space-time receptive fields of the LGN cells are shown before and after visual experience at the right. In this hypothesis, the X cell becomes more sustained with experience. The cortical neuron would be selective for motion that arose on the side of the X cell’s input location and moved in the direction of the transient input. If LGN cells underwent experience-dependent changes in their sustainedness or transience, direction-selectivity in V1 could be amplified. **b.** A direction-selective V1 cell is created by combining inputs selective for specific spatial locations in the cell’s receptive field and are stimulated by a moving stimulus in a precise temporal order. In this case, a direction-selective V1 cell could be created if the Y cell’s latency was reduced by experience, resulting in a short-latency LGN cell input (SL) at one location and a longer latency LGN cell input (LL) at a neighboring location. If LGN cells exhibited changes in latencies with experience, direction selectivity in V1 could be amplified.

We examined the sustainedness and transience of LGN neurons using counterphase sinusoidal gratings (0.08 cycles/degree; temporal frequency 1 Hz) presented at 6 different spatial phases. These stimuli, when shown at a phase that lined up with the receptive field center, flipped the stimulus provided to the center from ON to OFF each 0.5 seconds. We chose the spatial phase that elicited the strongest response (F1) for analysis. From the PSTH we found that the peak firing rate of the LGN neurons did not significantly change after exposure to the training stimuli (t(54)=−0.0893, p=0.9291) (**Figure 4a-e**). The maintained rate also did not significantly change after exposure to the training stimuli (t(54)=−1.5392, p=0.1296) (**Figure 4f**). The transience values for the naive LGN neurons (median=0.9008) changed very slightly with exposure to training stimuli (median=0.87455; t(54)=1.9932, p=0.0513), but not significantly (**Figure 4g**). In summary, 6 hours of visual experience had no significant effect on orientation or direction selectivity, or sustainedness or transience of LGN neurons. This suggests that there was no change in the availability of sustained or transient signals to cortex as a result of short-term experience.

**Figure 4.**
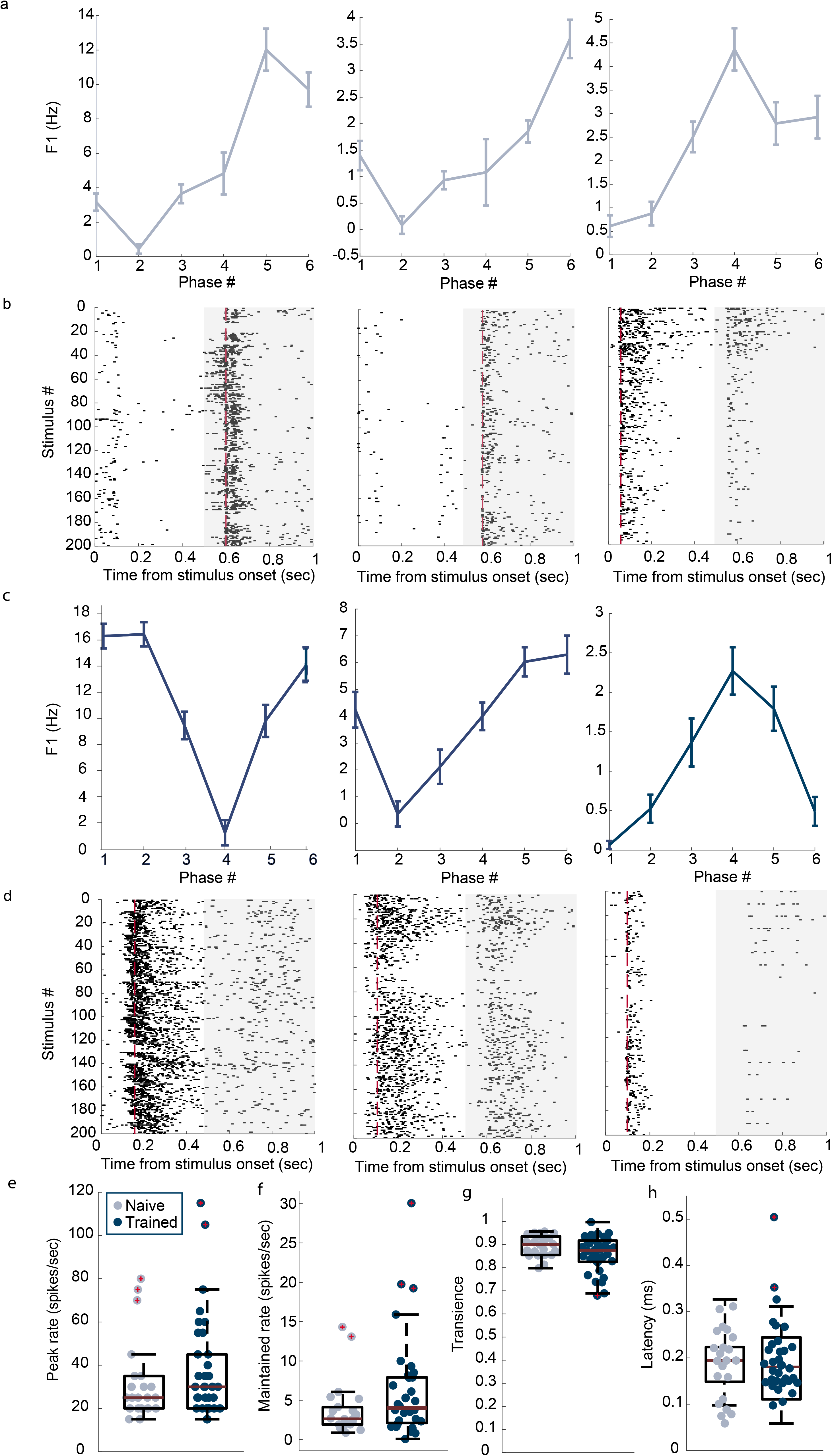
Spatial phase analysis and sustainedness and transience and latency of LGN cells. **a.** 3 example naïve LGN neuron responses (F1) to six different spatial phases. **b.** Raster plots from Figure 4a naïve LGN neurons responses to the spatial phase that elicited the strongest response. Responses from 200 trials were averaged for peristimulus time histograms (PSTH; 10-ms time bins) of cell responses. White background indicates one phase and grey background indicates new phase. From the PSTH of the response we computed peak response (red dashed line), maintained response, and transience. **c.** 3 example trained LGN neuron responses (F1) to six different spatial phases. **d.** Raster plots from Figure 4c trained LGN neurons responses to the spatial phase that elicited the strongest response. Conventions as in 4b. **e.** Boxplot of peak rate values for naïve (before) and trained (after) LGN cells. Central line indicates median, top and bottom of box indicate 25th and 75th percentiles, whiskers indicate minimum and maximum values not considered outliers, outliers indicated by red cross. There was no significant change in peak rate with training (t(54)=−0.0893, p=0.9291). **f.** Boxplot of maintained rate values for naïve (before) and trained (after) LGN cells. Conventions as in Figure 4e. There was no significant change in maintained rate with training (t(54)=−1.5392, p=0.1296)). **g**. Boxplot describing transience of cells before and after training. Conventions as in Figure 4e. There was no significant difference in transience with training (t(54)=1.9932, p=0.0513). **h.** Boxplot of cells’ response latencies before and after training. There was no significant change in response latency after training ((54)=−0.5325, p=0.5966).

### Latency

In our next set of analyses, we examined changes in response latency. According to the Reichardt model (Reichardt, 1961), maximum activation of a direction-selective cell occurs when its inputs are selective for specific spatial locations in the cell’s receptive field and are stimulated by a moving stimulus in a precise temporal order. In this case, when a stimulus moves in the cell’s preferred direction it activates the inputs with longer latencies first and progresses towards activation of shorter latencies, allowing for the subthreshold inputs to simultaneously arrive at the cell’s soma and summate, causing an action potential (**Figure 3b**). A stimulus moving in the opposite direction, therefore, would activate cells with the shortest latencies first and progress towards longer latencies and a subthreshold response by the postsynaptic cell. If we saw a change in response latencies of thalamic inputs with short-term experience, such as a sharpening of latency tuning, this might suggest that a precise arrangement of thalamic inputs onto cortical cells contributes to direction selectivity development (Reichardt and Poggio, 1976; Reichardt, 1987).

To test this model, we examined initial response latencies of naïve LGN cells and compared these to response latencies following exposure to training stimuli. We found that initial latencies did not significantly change with 6 hours of visual experience (t(54)=−0.5325, p=0.5966; **Figure 4h**). This suggests that short-term experience does not cause a change in response latencies of thalamocortical inputs.

### Spatial and temporal frequency

Although we found no change in response latencies, we were interested in assessing whether there were changes in spatial frequency tuning and temporal frequency tuning with short-term visual experience. The primary reason we co-varied spatial and temporal frequency with direction was to ensure that we assessed orientation and direction at each cell’s preferred spatial and temporal frequency, but this data allows us to examine spatial and temporal frequency tuning for stimulation in the preferred direction. We examined cells’ responses to six different spatial frequencies and six different temporal frequencies using moving sinusoidal gratings at each cell’s preferred direction (**Figure 5a,b**). We found no significant change in spatial frequency tuning with visual stimulation training (Kruskal-Wallis test, H(1)=0.0044, p=0.9470; **Figure 5c**).

**Figure 5.**
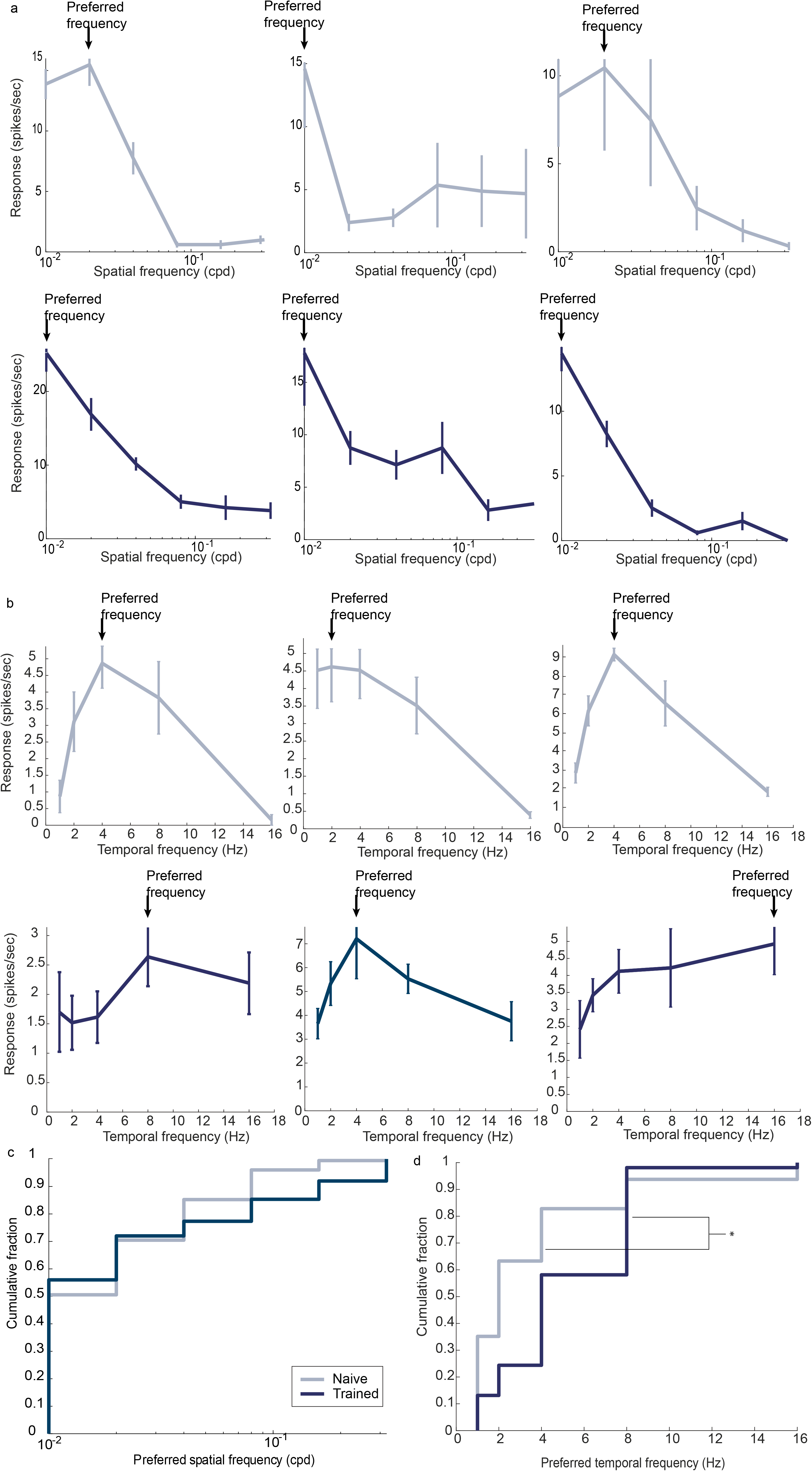
Spatial and temporal frequency analysis of LGN cells. **a**. Spatial frequency tuning curves of 3 representative LGN cells before 6 hours of visual stimulation training (top row) and after training (bottom row). Top and bottom tuning curves do not come from the same cell. Error bars indicate standard error. **b.** Temporal frequency tuning curves of 3 representative LGN cells before 6 hours of visual stimulation training (top row) and after training (bottom row). Conventions as in Figure 5a. **c.** Cumulative histogram of LGN cells preferred spatial frequency (cpd) before and after training. Preferred spatial frequencies did not significantly increase with training (Kruskal-Wallis test, H(1)=0.0044, p=0.9470). **d.** Cumulative histogram of LGN cells preferred temporal frequency (Hz) before and after training. Preferred temporal frequencies significantly increased with training (Kruskal-Wallis test, H(1)=32.7565, p=1.0446e-8).

We similarly examined cells’ responses to differing temporal frequencies. In previous recordings in visual cortex (Ritter et al., 2017), temporal frequency preferences increased over the duration of experiments for animals that were provided with 6-9 hours of experience with either drifting gratings or control animals that merely watched a gray screen for 6-9 hours. These increases in temporal frequency preferences occurred in all groups, including those that exhibited increases in direction selectivity (the animals that watched moving gratings) and those that did not exhibit increases in direction selectivity (the animals that watched a gray screen). Therefore, we expected to see an increase in temporal frequency preferences with the duration of the experiment. Indeed, when we examined cells’ responses to different temporal frequencies, we found a significant increase in the preferred temporal frequency of LGN neurons with training (Kruskal-Wallis test, H(1)=32.7565, p=1.0446e-8; **Figure 5d**). The mechanism of this process is unknown, but these increases in temporal frequency did not correlate with increases in direction selectivity in previous studies and rather were only correlated with the duration of the experiment (Ritter et al., 2017).

## Discussion

We assessed the impact of early visual experience on developing LGN neurons in visually naïve ferrets by characterizing the response properties of immature LGN neurons and the impact of moving visual stimuli. We found that naïve ferret LGN cells did not exhibit strong orientation or direction selectivity. Further, the low orientation and direction selectivity index values of ferret LGN neurons were not increased by 6 hours of exposure to a moving stimulus. Other receptive field properties that might influence inputs to a circuit that computes direction selectivity, such as the degree of sustainedness and transience or response latency, were also not altered by short-term experience. These data suggest that orientation and direction selectivity, and early changes in visual response properties, develop downstream from LGN. These results are consistent with the idea that V1 is the earliest stage in the ferret visual system that demonstrates orientation and direction selectivity, and that the development of mature, selective responses occurs entirely in cortex.

### Orientation and direction selectivity in LGN

Previous studies in mice and rabbit (but not squirrel: (Zaltsman et al., 2015)) have shown that there is a high percentage of orientation- and direction-selective cells in LGN (Swadlow and Weyand, 1985; Marshel et al., 2012; Piscopo et al., 2013; Scholl et al., 2013; Zhao et al., 2013; Suresh et al., 2016) and that, in mice, cortex receives orientation-selective inputs from LGN (Cruz-Martin et al., 2014; Kondo and Ohki, 2016; Sun et al., 2016). In contrast, we found that LGN cells in ferret are not strongly tuned for orientation and direction at the onset of visual experience, or with short-term visual experience. While our surveys were limited to the part of the visual field that was directly in front of the animal, cortical neurons that serve the same visual region exhibit substantial direction selectivity, and indeed, it is a property of a typical cell (Gilbert, 1977; Weliky et al., 1996; Clemens et al., 2012). Our findings imply that individual thalamocortical cells do not significantly contribute to direction selectivity in cortex. Had it been the case that LGN presynaptic inputs to V1 were orientation- and direction-selective, it could have been possible that LGN was performing the computation that contributes to cortical direction selectivity, and that V1 was inheriting direction selectivity.

### It is unlikely that LGN cells contribute to the rapid emergence of direction selectivity

Before this study, what roles could LGN cells have conceivably played in the rapid development of direction selectivity in ferret that occurs following 3-9 hours of visual stimulation with moving stimuli? Li/Van Hooser et al. (2008) showed that direction selectivity develops only for the orientation columns that are stimulated – for example, if the animal observed vertical stimuli moving left and right, then direction selectivity developed in the vertical orientation columns but not in the horizontal orientation columns. Therefore, if changes in LGN receptive fields could have contributed exclusively to this increase in direction selectivity, one would have to imagine that it would be through increases in direction selectivity in individual LGN cells that somehow projected to the vertical orientation columns. This outcome was unlikely, as it is at odds with the prevailing “spot detector” view of most LGN receptive fields. Nevertheless, the young brain is highly plastic and it was prudent to check for this possible contribution. Our results here do not provide any evidence in favor of an increase in orientation or direction selectivity for LGN cells, so it is highly unlikely that such an increase is responsible for the rapid increases in cortical direction selectivity.

A more likely possibility was that changes in both cortex and LGN could have contributed to the rapid increases in cortical direction selectivity with experience. An optogenetic study that provided direct stimulation of the cortical surface for 9 hours showed that activity provided to cortex alone, without direct stimulation of the LGN, was sufficient to increase direction selectivity in the naïve ferret cortex (Roy et al., 2016). These experiments showed that cortical stimulation and resulting changes were enough, in principle, to cause increased direction selectivity. However, these experiments did not tell us that plasticity in cortex was the only process that contributes to the increases in direction selectivity with acute visual experience. It was possible that concomitant experience-dependent changes in LGN receptive fields also contributed to experience-dependent changes that were observed in cortical cells.

In this paper, we outlined how changes in the available LGN cell latencies or increases in sustainedness or transience could have contributed to increased direction selectivity by providing cortex with more reliable information about stimulus direction. Changes of this type would not have been enough, by themselves, to explain the increases in direction selectivity in ferret: presumably, all LGN cells that are activated by moving gratings would have undergone these changes, yet Li/Van Hooser et al. (2008) only observed changes for particular orientation columns. Clearly, changes in cortex that were specific to the orientation columns stimulated would also have been needed to take advantage of the increase in information that might have been provided by LGN cells, but it would have allowed for a dual role of LGN and cortex in the development of direction selectivity.

Instead, we found no evidence for short-term changes in LGN receptive field latencies or sustainedness/transience, and no overall evidence of changes in LGN that might have contributed, even in part, to the rapid emergence of direction selectivity in cortex. Given a) the orientation specificity of the emergence of direction selectivity (Li et al., 2008) and that orientation selectivity is first found in cortex (Hubel and Wiesel, 1959), b) the fact that direct optogenetic stimulation of the cortex causes emergence of direction selectivity (Roy et al., 2016), and c) we find no evidence of changes in LGN receptive fields with short term experience (this paper), we conclude that the rapid emergence of direction selectivity in ferret visual cortex involves only processes that are taking place in the cortex. These changes could include modifications of LGN to V1 cell connections in cortex, modification of intracortical connections, and also include, in part, changes to cortical neuronal excitability (Roy et al., 2020).

### Rapid development vs. normal development

Under normal conditions, the experience-dependent development of direction selectivity in carnivore visual cortex takes place over about two weeks after eye opening (Li et al., 2006). During this time, LGN receptive fields do become smaller and exhibit shorter latencies (Tavazoie and Reid, 2000), temporal frequency preferences increase (Cai et al., 1997), and the latencies of the fastest inputs to visual cortical neurons also decrease (Roy et al., 2020). The results here show that these changes in LGN receptive fields are not necessary for increases in cortical direction selectivity.

This result has important implications for the circuit mechanisms that underlie direction selectivity in V1. Many models that are influenced by the Reichardt (Reichardt, 1961) model rely on precise delays of feed-forward inputs at different locations (Saul and Humphrey, 1990; Feidler et al., 1997; Van Hooser et al., 2014), but direction selectivity can be formed acutely in cortex well before the final latencies of LGN cells are established. Because direction selectivity persists despite changes in the latencies of the input cells, these results provide indirect support for models of direction selectivity that do not require precisely-timed excitatory input.

One class of such models rely on feed-forward inhibition that modifies feed-forward excitation. This feed-forward inhibition might arise in an asymmetric manner via lateral cortical connections, such that more inhibition is triggered by stimulation on the null side, as in the rodent and lagomorph retina (Barlow and Levick, 1965; Euler et al., 2014). A study that used optogenetic stimulation uncovered asymmetric null-side inhibition onto layer 2/3 neurons in ferret visual cortex (Scholl et al., 2019), and it’s possible that this configuration is found in layer 4 as well. However, intracellular studies by Priebe and Ferster (2005) found no evidence for strong null-side inhibition in layer 4 neurons in cat. Instead, disynaptic feed-forward inhibition that arrives at particular phases of stimulation may confer direction selectivity (Priebe and Ferster, 2005; Freeman, 2021). A mechanism like this might be robust to refinements in LGN cell latency, as the disynaptic delay from feed-forward interneurons to cortical excitatory neurons would be relatively constant even as the latencies of the feed-forward inputs changed.

Another feed-forward model that does not require such exquisite timing of LGN inputs is the sustained/transient convergence model (Marr and Ullman, 1981; Lien and Scanziani, 2018). If inputs at one side of the receptive field are transient and inputs at the other side of the receptive field are sustained, then the temporal summation of these two inputs could still produce a direction-selective signal, even if the absolute latencies of these inputs changed over development.

## Acknowledgements

This work was funded by NIH EY022122. We thank David Landesman for initial contributions to the chamber prototype. We thank members of the Van Hooser lab for comments.

## Notes

Conflicts of interest: The authors declare no competing financial interests or other competing interests.

### Competing Interest Statement

The authors have declared no competing interest.

